# DNA Recognition and Induced Genome Modification by a Hydroxymethyl-γ Tail-Clamp Peptide Nucleic Acid

**DOI:** 10.1101/2023.07.08.548209

**Authors:** Stanley N. Oyaghire, Elias Quijano, J. Dinithi. R. Perera, Hanna K. Mandl, W. Mark Saltzman, Raman Bahal, Peter M. Glazer

## Abstract

Peptide nucleic acids (PNA) can target and stimulate recombination reactions in genomic DNA. We have reported that gamma (γ)-PNA oligomers possessing the diethylene glycol γ-substituent show improved efficacy over unmodified PNAs in stimulating recombination-induced gene modification. However, this structural modification poses a challenge because of the inherent racemization risk in *O*-alkylation of the precursory serine side chain. To circumvent this risk and improve γPNA accessibility, we explore the utility of γPNA oligomers possessing the hydroxymethyl-γ moiety for gene editing applications. We demonstrate that a γPNA oligomer possessing the hydroxymethyl modification, despite weaker preorganization, retains the ability to form a hybrid with the double-stranded DNA target of comparable stability and with higher affinity to that of the diethylene glycol-γPNA. When formulated into poly(lactic-co-glycolic acid) nanoparticles, the hydroxymethyl-γPNA stimulates higher frequencies (≥ 1.5-fold) of gene modification than the diethylene glycol γPNA in mouse bone marrow cells.

## INTRODUCTION

Peptide nucleic acids (PNAs) that bind to and stimulate recombination events in genomic DNA are useful reagents for genome editing^1–8^, and avoid or minimize the emerging limitations of nuclease-based^9–11^ reagents (like CRISPR-Cas9) such as genotoxicity^12–16^ and immunogenicity^17, 18^. Developed as nucleic acid mimics and ligands^19, 20^, PNA oligomers bind DNA targets with high affinity, activating—in the context of genomic DNA—endogenous DNA repair pathways that mediate incorporation of co-supplied donor DNA oligomers carrying the intended sequence changes at the target (homologous) site^1^. Further, PNAs’ binding specificity^21^ restricts repair activation and resultant gene modification to genomic regions at/proximal to the PNA binding site^22–24^, allowing for gene editing to proceed with low off-target effects^5–8^. Importantly, when encapsulated in and delivered by poly(lactic-co-glycolic acid) (PLGA) nanoparticles (NP)^25^, PNA/donor oligomers mediate gene correction at frequencies sufficient to reverse disease phenotypes in mice^6–8^.

The efficacy of PNA oligomers for gene editing is connected to their affinity for complementary DNA targets that can exist transiently as accessible single strands, during DNA replication and gene transcription, for example, but predominantly as compact duplex structures^26^. For example, we demonstrated that the editing efficacy of a bisPNA oligomer^27^—which contains domains to recognize the Watson-Crick and Hoogsteen faces of a DNA target—was enhanced five-fold by extending the binding domain on the Watson-Crick face, as in a tail-clamp (tc) PNA design^28^, due to improved DNA invasion/binding^5^. Also, we recently reported that the incorporation of γPNA^29, 30^ monomers—derivatives that improve affinity and specificity of composite oligomers^29^ and their ability to invade B-DNA^31^—in a tcPNA oligomer increased editing frequency two-fold over the unmodified tcPNA^5^. The γPNA modification has been shown to drive conformational selection in PNA oligomers by inducing a network of steric clashes and nucleobase stacking interactions^30^, with the resultant helical sense determined by the stereochemistry at the PNA γ position^32^. Specifically, the *R*-configuration in γPNA monomers induces PNA oligomers to adopt a right-handed helical conformation as single, unbound strands^30^—the same orientation enforced on them upon binding to complementary DNA/RNA targets^33^. This pre-organization reduces the entropic cost of DNA binding, an effect that accelerates hybridization to single-stranded (ss) targets^29^, and enhances strand invasion of duplex targets^31^.

The chemical moiety at the γ position primarily depends on the nature of the precursory amino acid side chain^30, 34–40^, although additional synthetic derivatization is possible^29, 41–43^. To date, however, only γPNA monomers possessing the diethylene glycol (also called minipeg, mp ^29^) γ-substituent have been tested for gene editing applications and shown to enhance editing frequencies over unmodified PNAs^7, 8^. Minipeg-modified γPNAs (^mp^γPNA) retain all the biophysical features and improved binding properties of γPNA oligomers outlined above and possess enhanced aqueous solubility due to the hydrophilic mp moiety^29^. However, synthesis of the requisite monomers is challenging, primarily because of the racemization risk inherent in the crucial *O*-alkylation step to incorporate the mp unit unto the serine side chain^29^. In this regard, a variety of safeguards have been utilized to obtain optically pure monomers—ranging from careful control^29^ of reaction temperature and time, and reagent addition order—to *in situ*^29^ or precursory^44^ derivatization of serine to minimize the acidity of the alpha proton. However, these precautions do not obviate the need for further analytical validation of optical purity^29, 44^, a parameter especially salient in gene editing, since monomers with even minute (5%) racemates significantly compromise DNA affinity^45^.

We examined here whether γPNA oligomers possessing the hydroxymethyl γ-substituent would be effective reagents for gene editing. As this moiety is directly accessible from the serine side chain (hence ^ser^γPNA), monomer synthesis circumvents the stereochemical contamination risk inherent in *O*-alkylation, simplifying the synthetic procedure considerably, as has been reported^30, 39, 40^. We synthesized a ^ser^γPNA (from commercially available monomers) and compared its helical organization, DNA binding properties, and gene editing frequencies to a known isosequential ^mp^γPNA^7^. Although possessing the less elaborate γ-substituent, ^ser^γPNA preserves (but relaxes) the helical organization inherent in ^mp^γPNA. Surprisingly, the thermodynamic stabilities of hybrids formed with a double-stranded DNA target are comparable for both ^ser^γPNA and ^mp^γPNA, in both low and high-salt buffers, even with the weaker preorganization in the former. Importantly, we show that ^ser^γPNA induces higher (≥ 1.5-fold) editing frequencies with two different donor DNA oligomers, relative to ^mp^γPNA, in bone marrow cells from two different transgenic animals.

## RESULTS

### Rationale and γPNA Design

We previously reported that a tcPNA oligomer (PNA, Chart 1) designed to bind 70 bp upstream of the position of a β-thalassemia-associated mutation was able to stimulate recombination of a donor DNA oligomer with the target homologous region in genomic DNA^7^. The tcPNA design has binding domains that recognize the Watson-Crick and Hoogsteen faces of a purine-rich sequence of DNA target, with an extended recognition sequence on the Watson-Crick domain^28^. Modification of PNA with γPNA residues bearing the diethylene glycol γ-substituent on the Watson-Crick domain (^mp^γPNA, Chart 1) improved editing frequencies two-to three-fold *ex vivo* due to improved DNA binding. Nanoparticle-mediated administration of this PNA reversed thalassemic phenotypes in mouse models *in vivo*^7, 8^. To obviate the need for the elaborate chemical syntheses required to install the diethylene glycol unit in the precursor serine side chain^29, 44^, we explored whether a tcPNA bearing just hydroxymethyl γ-substituents, themselves directly derivable from serine (^ser^γPNA, Chart 1), would retain the biophysical and DNA hybridization properties of ^mp^γPNA, and remain superior for gene editing relative to PNA.

**Chart 1:**
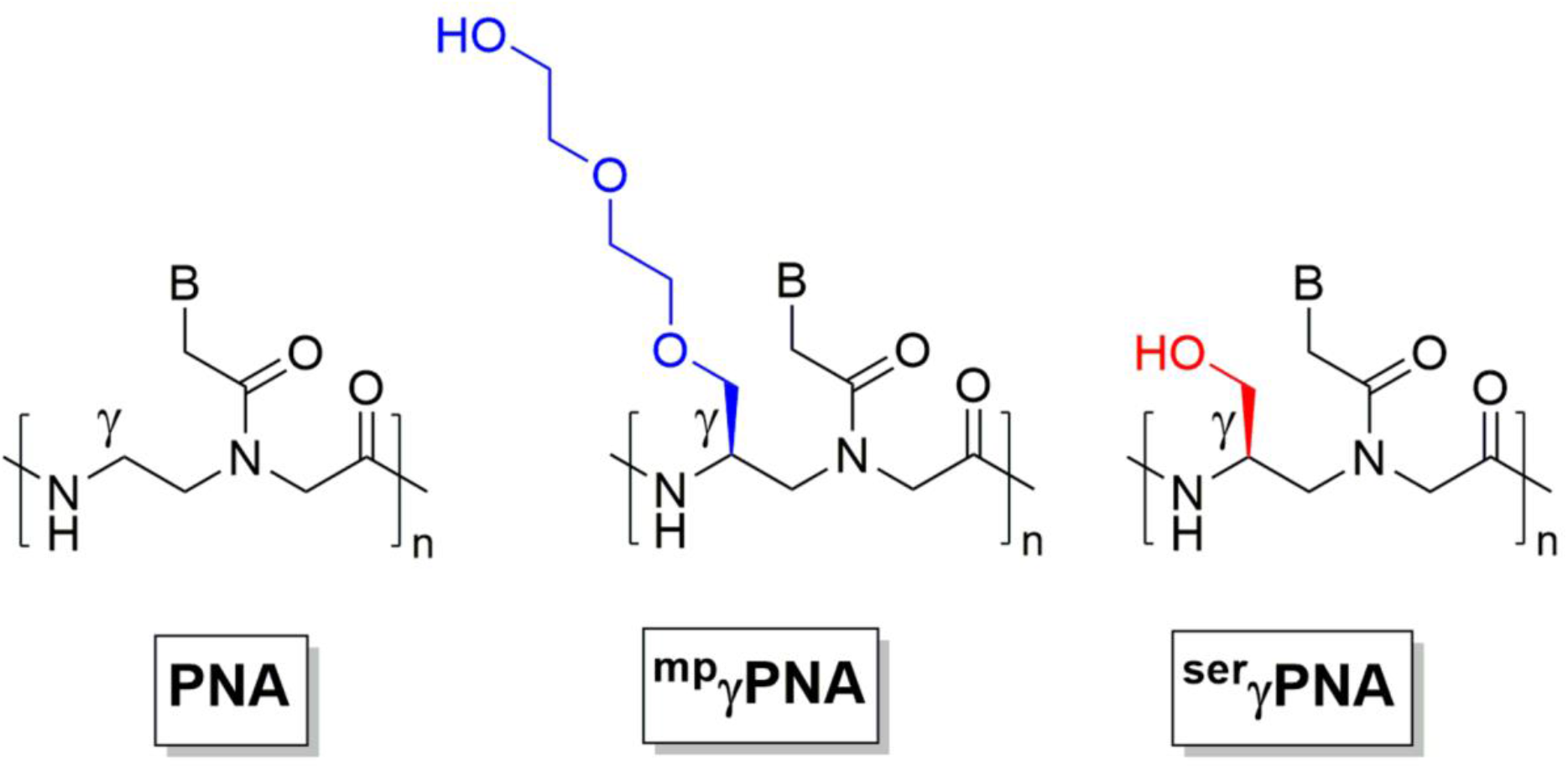
Chemical structures of PNA, ^mp^γPNA, and ^ser^γPNA monomer units.

### Helical Organization of PNA and γPNAs

We first sought to examine the effects of ^mp^γ-or ^ser^γ-PNA residues (Charts 1) on the global helical structure of the composite tcPNA oligomers using circular dichroism (CD) experiments, which have previously been utilized for characterization of γPNA helical conformation^29, 30, 46^. Our experiments show that samples containing equimolar amounts of the ^ser^γPNA or ^mp^γPNA oligomers display local minimum and maximum at 240-and 267 nm, respectively (in addition to the other peaks), both of which are absent in the unmodified PNA oligomer (Figure 1). The position (λ) and orientation (minima/maxima) of this specific exciton coupling pattern are consistent with the adoption of a right-handed helical structure^47^. However, the more pronounced 240 nm minimum observed for ^mp^γPNA suggests that it is more helically pre-organized^29^ than ^ser^γPNA. It is likely that the steric clashes in the γPNA backbone, which potentiate conformational selection in the oligomer, will be greater with the larger diethylene glycol (^mp^γPNA) than with hydroxymethyl (^ser^γPNA) as the γ-substituent, as suggested^48^ and demonstrated^32^ by Ly and coworkers. While helical organization occurs to varying degrees, it is concentration independent for both, as demonstrated by the linear correlation between signal amplitude and oligomer concentration (Figure S1), suggesting that the observed effects are intrinsic to the molecules themselves, as previously observed^30^, and are uninfluenced by aggregation or intermolecular complexation in solution. Interestingly, regardless of the absence or degree of helical organization in the PNA/γPNA oligomers, their hybrid triplexes with a complementary ssDNA target have similar helical structures (Figure 2), indicating that γ-modifications of either kind do not affect the conformation of the thermodynamic product from the binding reaction. The same trend was observed for PNA/γPNA binding in the context of a dsDNA target (Figure S2).

**Figure 1:**
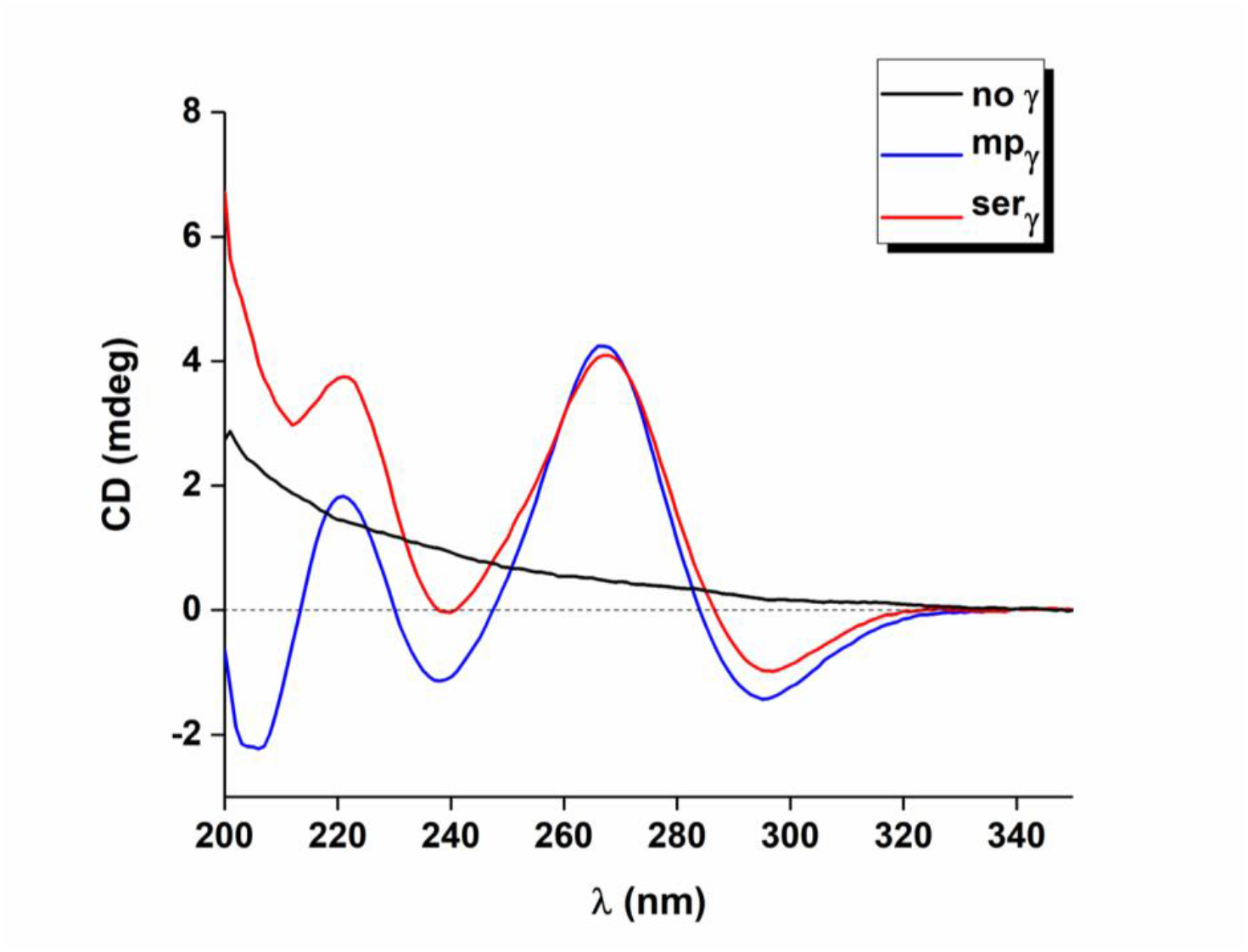
CD spectra for PNA (regular PNA, black), ^mp^γPNA (^mp^γ, blue), and ^ser^γPNA (^ser^γ, red). All samples contained 20 µM of the respective oligomer in 10 mM NaPi buffer. Spectra were recorded at 37 °C.

**Figure 2:**
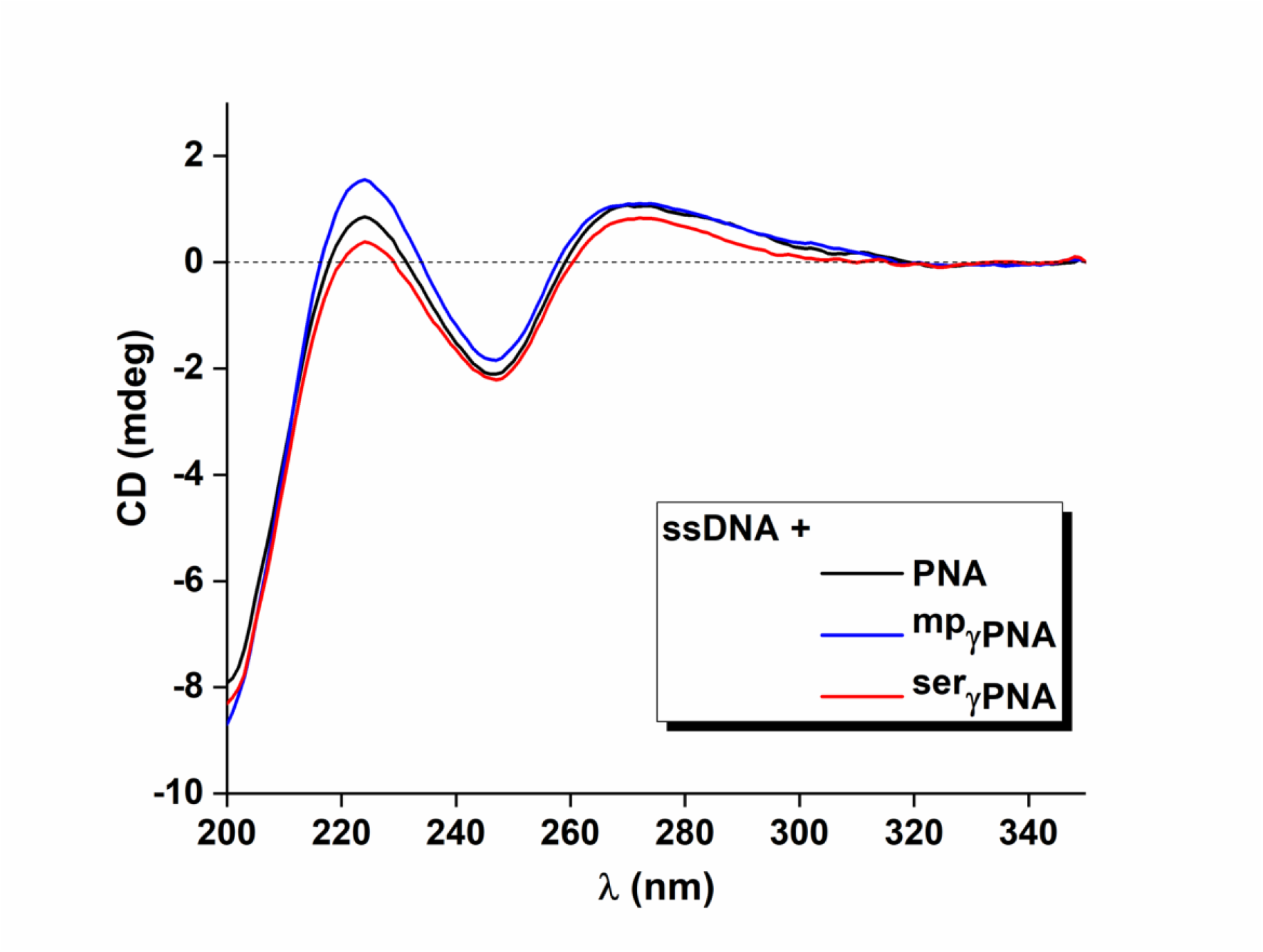
CD spectra for hybrid triplexes containing ssDNA and PNA (black), ^mp^γPNA (blue), or ^ser^γPNA (red). All samples contained 1 µM ssDNA and 2 µM PNA/γPNA in 10 mM NaPi buffer. Spectra were recorded at 37 °C.

### Thermal Stability of PNA/γPNA-DNA Hybrids

We next sought to characterize the thermal stabilities of the hybrid complexes by recording UV absorbance at 265 nm with increasing temperature. Initial analyses on hybrids formed with the ssDNA target yielded incomplete, noncooperative melting transitions, presumably due to high melting temperatures that preclude complete melting (Figure S3). We therefore present data (Figure 3) for hybrids formed with a double-stranded DNA target (dsDNA) preassembled by annealing ssDNA with the PNA and then with its complement. The dsDNA-PNA/γPNA hybrids were assembled under conditions where complete dsDNA binding was observed by gel shift (Figure 4), enabling unambiguous attribution of the melting transitions to dissociation of the hybrid complex rather than to contributions from partially PNA-bound and free dsDNA.

**Figure 3:**
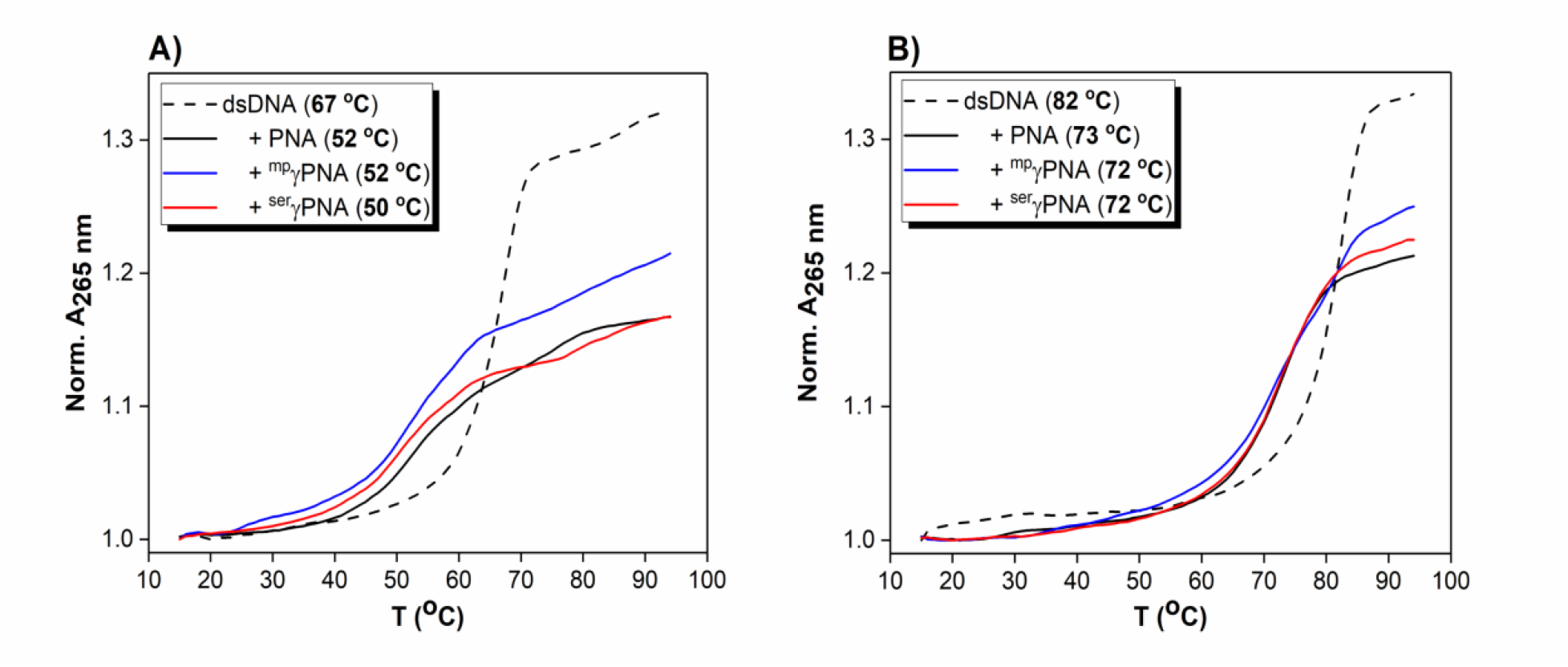
UV-thermal denaturation analyses on a 90mer dsDNA alone (broken black) or in combination with PNA (solid black), ^mp^γPNA (blue), or ^ser^γPNA (solid red). All samples contained 2.5 µM dsDNA and 5 µM PNA/γPNA in 10 mM NaPi (A) or 100 mM NaPi (B). All samples were annealed as described in Methods and melting Temperature (Tm) values are presented in parentheses in the legend.

**Figure 4:**
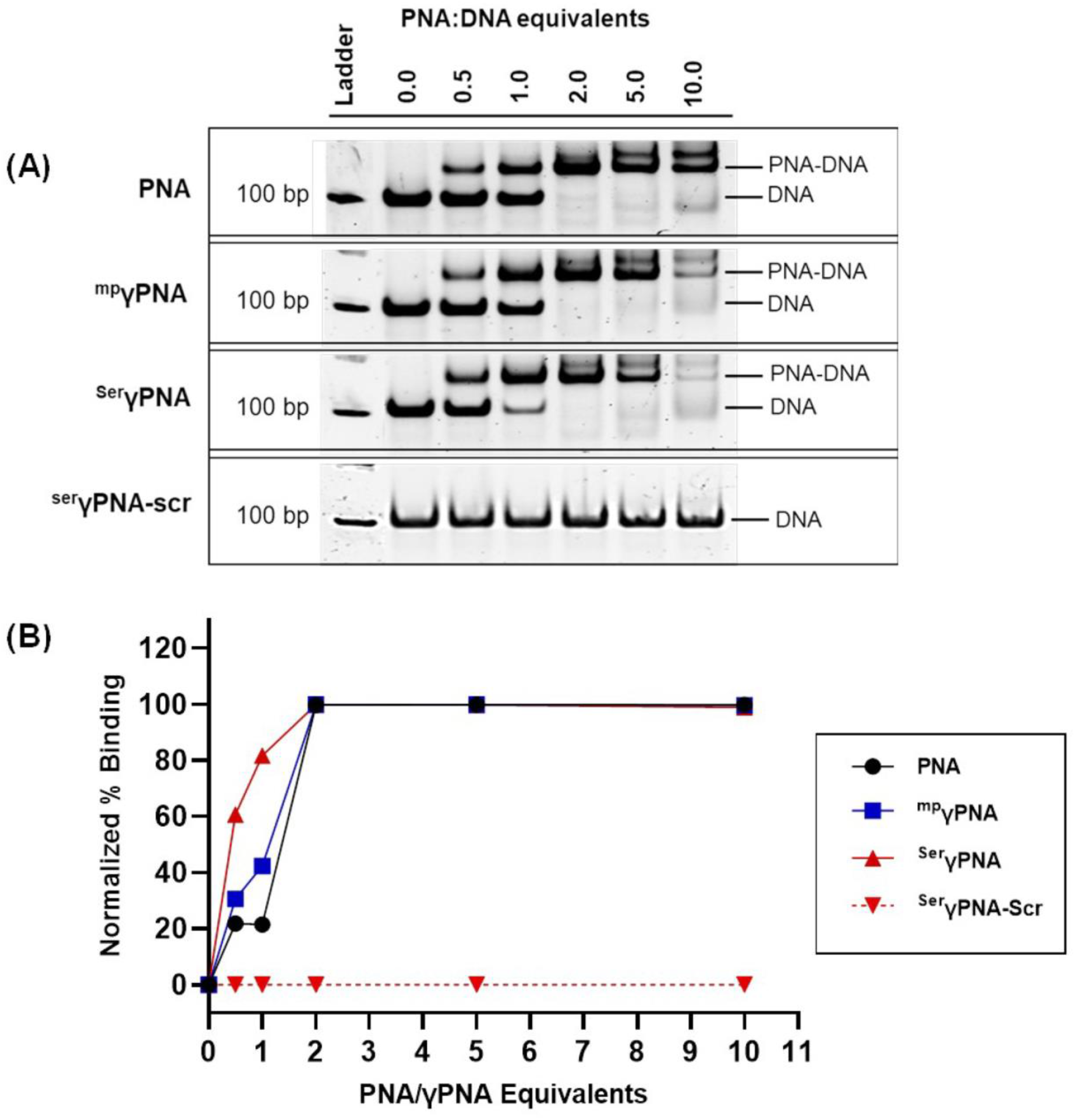
Titration of 100mer dsDNA target with increasing equivalents of PNA, ^mp^γPNA, ^ser^γPNA, or ^ser^γPNA-scr. Each sample contained 50 nM DNA in 100 mM NaPi buffer, was annealed as in Methods, and was run on an 8 % PAGE gel (A). The band intensities for the PNA-DNA hybrids were measured on ImageJ and normalized to the 0 eqv condition (B).

We observe that the hybrid complexes are less thermally stable than the free dsDNA (-ΔT = 15 -17 °C, Figure 3A and Table 2) in 10 mM NaPi, consistent with a binding model where PNA/γPNA hybridization to the target strand is accompanied by displacement of the homologous region on the non-target DNA strand. In this model, the hyperchromicity accompanying melting likely reflects the release of the non-target strand from the PNA-bound target strand, since ssDNA-PNA/γPNA hybrids show incomplete transitions. Further, similar analyses at 100 mM NaPi show that although the same binding-induced duplex destabilization occurs, the melting temperatures for the hybrids are higher (Figure 3B and Table 2) than that at 10 mM Na Pi, presumably because the DNA-DNA base pairs flanking the PNA-bound region are made more stable at the higher salt concentration.

We also performed van’t Hoff analyses of the melting profiles using the protocols outlined by Marky and Breslauer^49^. We observe complex formation as an exergonic process for all PNA/γPNAs at 10 and 100 mM NaPi [-ΔG = 34-35 kcal mol^-1^ and 41-45 kcal mol^-1^, respectively (Table 2)]. The complexes are more stable at 100 mM NaPi, an unsurprising trend in our model, since the DNA-DNA base pairs on either side of the PNA-DNA hybrid, which remain unperturbed by PNA recognition, would be more stable under this condition. There are also expected and surprising trends from the data: for example, -ΔS is least for the ^mp^γPNA-DNA hybrid, consistent with greater preorganization in the PNA; while -ΔH is smallest for the same PNA, contrary to data showing a strong enthalpic effect of ^mp^γ modification^29^.

### DNA Invasion by PNA/γPNAs

To evaluate the relative binding affinities of the PNA/γPNAs for strand invasion towards a duplex DNA target, we titrated increasing equivalents of the respective PNAs into 50 nM target dsDNA (100mer) in 100 mM NaPi buffer. Compared to diethylene glycol-γPNA and classic-PNA, the hydroxymethyl-γPNA showed superior strand invasion and binding to dsDNA (Figure 4).

### Nanoparticle Formulation and Characterization

Previously, we demonstrated that nanoparticles (NPs) made of poly(lactic-co-glycolic acid) (PLGA) could effectively encapsulate and deliver PNA and donor DNA to correct mutations underlying β-thalassemia *in vivo* in adult^7^ and fetal mice^8^. Using a similar approach, we synthesized PLGA NPs encapsulating ^mp^γPNA or ^ser^γPNA along with donor DNA to correct or introduce the same IVS2-654 mutation^7, 8^. The physiochemical characteristics of these NP formulations did not differ significantly between both γPNAs (Figure 5). Average NP diameters were 270 nm for ^mp^γPNA and 290 nm for ^ser^γPNA, as measured by dynamic light scattering (DLS) (Figure 5A). Likewise, NP surface charge was similar between both formulations, at -19 mV for ^mp^γPNA and -24 mV for ^ser^γPNA NPs (Figure 5B). Similarly, average total nucleic acid loading did not differ significantly (Figure 5C). Despite differences in chemical modification, both PNAs displayed similar release rates, with greater than 50% of total nucleic acid content being released for each formulation after 72 hours (Figure 5D).

**Figure 5:**
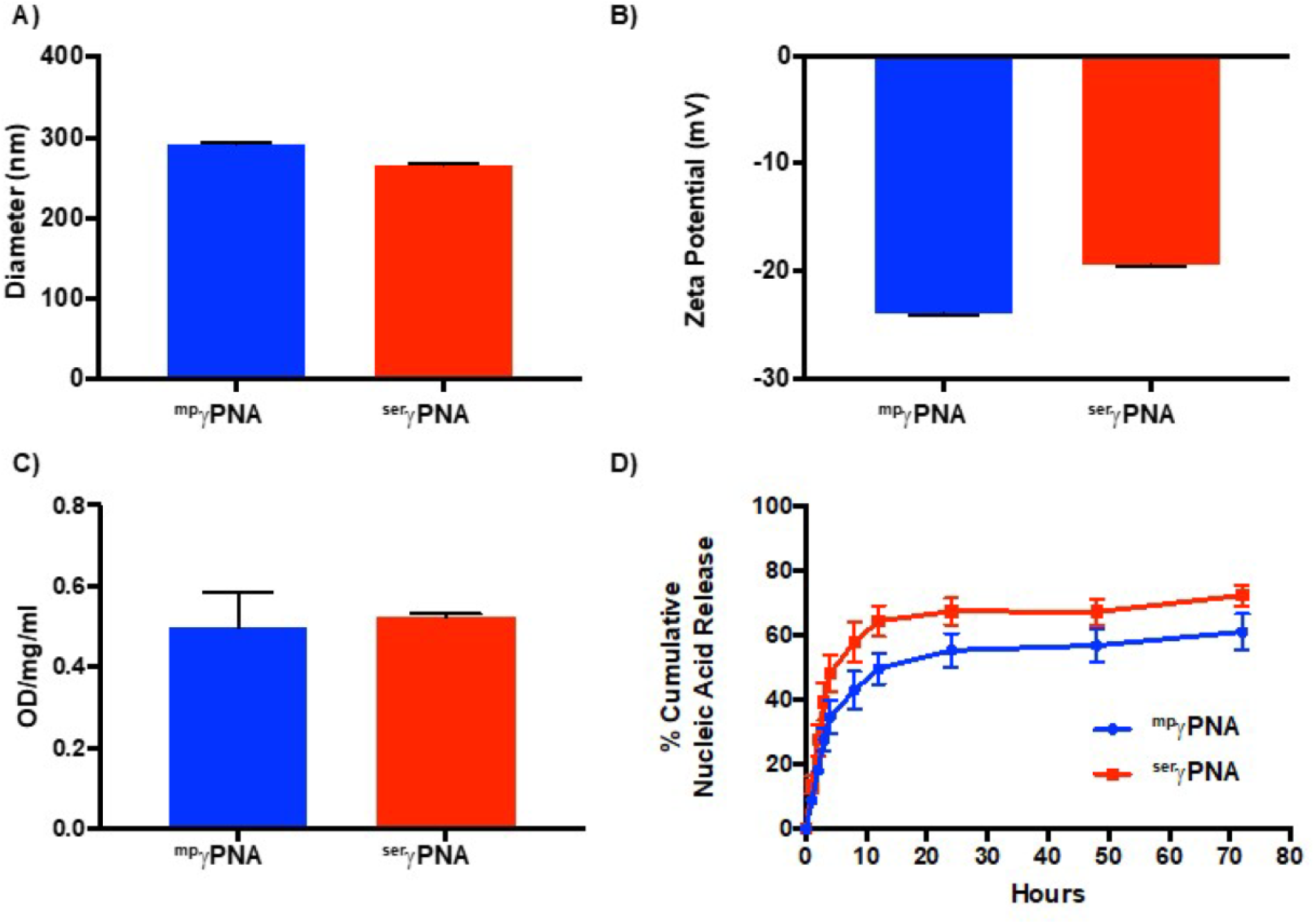
Nanoparticle characterization. (A) NP diameter as measured by dynamic light scattering (DLS). (B) NP surface charge as measured by zeta potential. (C) Total nucleic acid loading (PNA + Donor DNA) in PLGA NPs. (D) Release of nucleic acids from PLGA NPs.

### Correction of the β-thalassemia Mutation in Primary HSPCs

Given the physiochemical similarities between the NPs, we hypothesized that the superior hybridization and duplex invasion properties of ^ser^γPNA would result in higher levels of gene editing relative to ^mp^γPNA. To test this hypothesis, we started by designing a donor DNA to introduce the IVS2-654 mutation. NPs were subsequently formulated to encapsulate this ‘mutating’ donor DNA and ^mp^γPNA or ^ser^γPNA. Primary bone marrow cells from mice with a wild-type human β-globin transgene were isolated and treated with 2 mg ml^-1^ of PLGA NPs. We found (Figure 6) that NPs encapsulating ^ser^γPNA achieved significantly higher levels of genome modification, as measured by a droplet digital PCR (ddPCR) assay previously optimized to detect the wild-type and mutant alleles in genomic DNA^8^.

**Figure 6:**
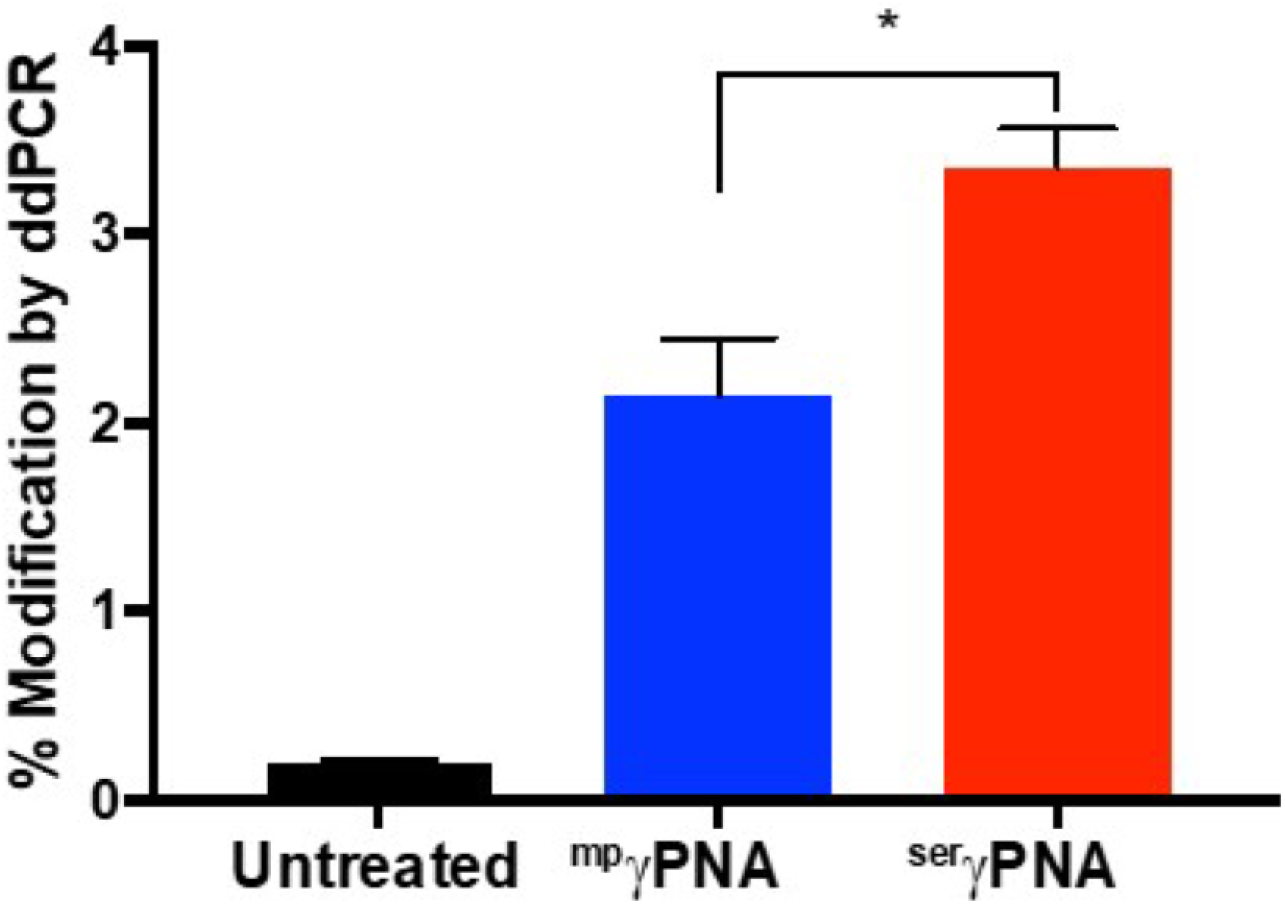
Modification of primary bone marrow cells with the β-thalassemia-causing mutation at IVS2-654. BMCs were left untreated (black) or were treated with NPs containing ^mp^γPNA (blue) or ^ser^γPNA (red).

We next sought to determine whether ^ser^γPNA could induce gene correction in bone marrow from a mouse model of β-thalassemia with the IVS2-654 mutation. As before, NPs were formulated with ^mp^γPNA or ^ser^γPNA but this time with a ‘correcting’ donor DNA. Primary bone marrow cells from these mice were treated with 2 mg ml^-1^ of PLGA NPs. Again, gene correction of the underlying β-thalassemia mutation was significantly higher with ^ser^γPNA (Figure 7), despite similarities in all the measured physiochemical properties of the NPs themselves. We also compared editing of ^ser^γPNA with ^ser^γPNA-scr and confirmed minimal editing with the scrambled oligomer compared to the targeting PNA (Figure S4). Taken together, these experiments show the superiority of ^ser^γPNA compared to the ^mp^γPNA in inducing genome modification, as observed in two distinct primary cell lines with two distinct donor DNA sequences.

**Figure 7:**
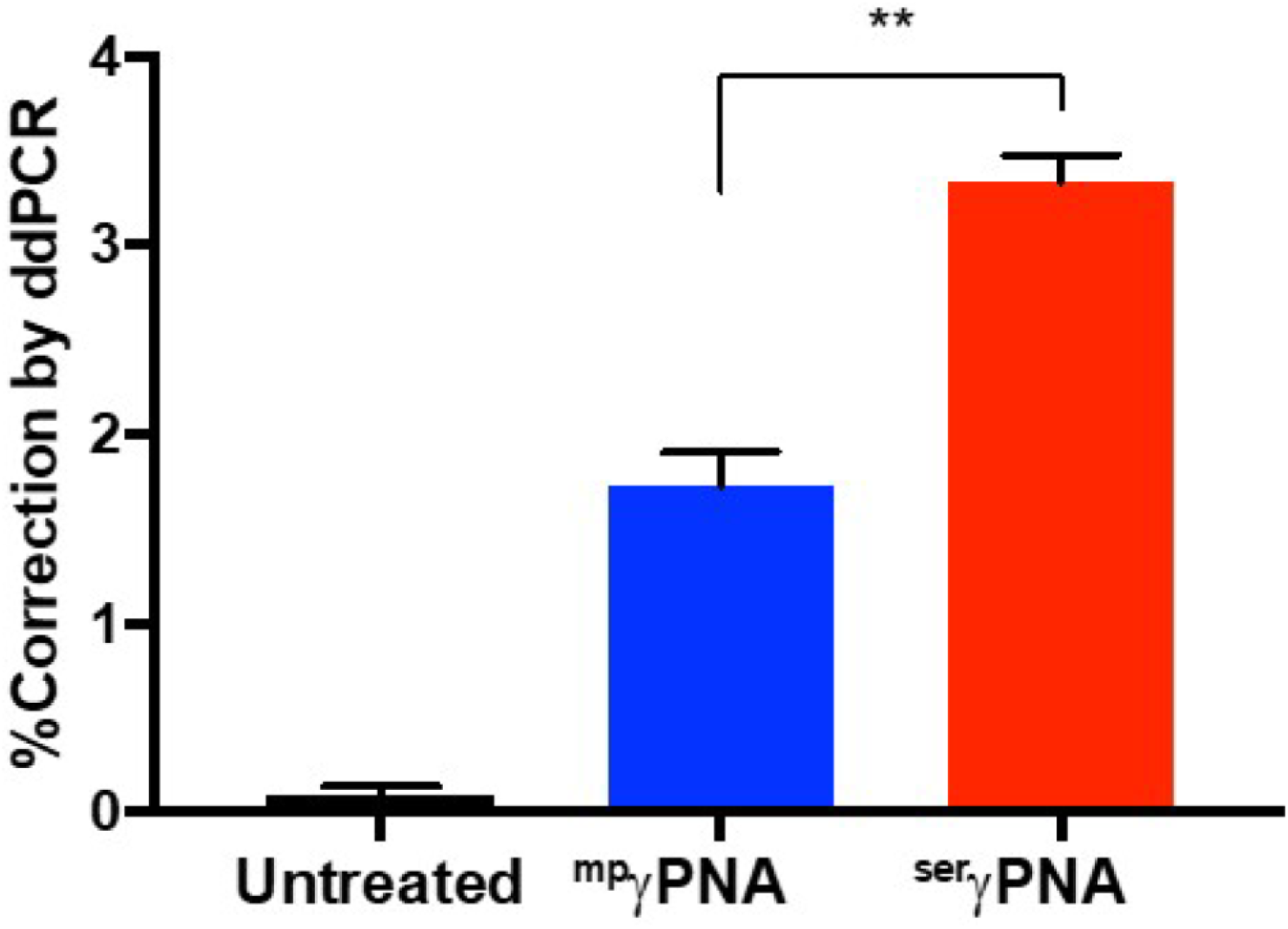
Correction of the β-thalassemia causing mutation in primary bone marrow cells. BMCs were left untreated (black) or were treated with NPs containing ^mp^γPNA (blue) or ^ser^γPNA (red).

## DISCUSSION

We present data on the biophysical and DNA-binding properties of a tcPNA oligomer possessing a hydroxymethyl γ(^ser^γ)-substituent and compare/contrast these with measurements for an isosequential oligomer with the mp-γ modification (^mp^γ). Our results indicate that, as unbound molecules, both oligomers adopt a right-handed helical conformation, as predicted from the configuration of the γ-stereogenic center^29, 30^. Helical organization is more pronounced for the ^mp^γPNA than the ^ser^γPNA, an observation predictable from the relative sizes of the chemical moieties at the γ-position^32^ (diethylene glycol and hydroxymethyl, respectively). DNA recognition in the context of a duplex target yields hybrids of comparable stabilities for both ^ser^γPNA and ^mp^γPNA, and we observe strand invasion of hydroxymethyl-γPNA to be superior to that of diethylene glycol-γPNA.

Because the ^mp^γPNA examined in this work was previously reported to mediate genotypic and phenotypic correction in β-thalassemic mice^7, 8^, we evaluated the efficacy of ^ser^γPNA for genome modification in bone marrow cells from transgenic mice harboring the human β-globin gene, and possessing a wild-type or mutant genotype at the relevant thalassemia-associated locus. In both formats, ^ser^γPNA induced higher gene editing frequencies than ^mp^γPNA, a trend not attributable to any differences in the physicochemical properties of the PLGA NPs synthesized to encapsulate and deliver the respective γPNA/donor DNA reagents. Taken together, our results show that the hydroxymethyl-γ modification, which is directly accessible via the serine side-chain and circumvents the synthetic limitations and racemization risks inherent in elaborate modification at the γ-position^29, 44^ (as in ^mp^γ), can be used as an alternative to the more specialized diethylene glycol-γ modification, yielding composite γPNA oligomers that show comparable DNA recognition properties and gene modification frequencies.

While they have been repurposed here, for the first time, as viable alternative/improved co-reagents for mediating PNA-induced gene modification, ^ser^γ-modified PNAs are not new in the literature of PNA-based nucleic acid ligands. In their seminal work describing the effects of PNA backbone γ-substitutions on the global oligomer structure, Ly and coworkers^30^ demonstrated that incorporation of hydroxymethyl γ-residues in a PNA oligomer introduced defined steric clashes in the PNA backbone that initiated a unidirectional helical preorganization^30^. They further showed that complete or partial modification of the PNA oligomer with this substituent, as with other moieties^29, 43, 48, 50^ introduced at the same position, resulted in improved binding to target DNA/RNA and enhanced selectivity against non-target strands compared to classic PNAs ^30^. Romanelli and coworkers subsequently showed that PNA oligomers sparingly modified with ^ser^γ residues effectively perturbed Sp1 recognition of a duplex DNA target modelled after its binding sequence in cognate promoters^40^. Virta and coworkers have also introduced these modifications into a PNA oligomer designed to target a model microRNA hairpin structure^39^. Our work extends these prior studies by demonstrating the superiority of the ^ser^γ-modified PNAs for binding to duplex DNA and for gene editing in primary bone marrow cells.

The utility of ^ser^γPNA in gene editing applications raises the question of what other existing^34–38, 41, 42^ or novel γ modifications may be repurposed or developed for similar ends, especially if they are more synthetically and/or commercially accessible than existing reagents. While we consider this question to be an interesting area for future study, we believe that at least two parameters should guide explorations on this theme: (1) hydrophilicity of the γ-substituents in order to preserve the aqueous solubility of the ^mp^γ-modified PNA reagents; and (2) substituent size in order to retain the steric clashes that initiate conformational selection in the oligomer. For the latter, careful tuning will be required to ensure that conformational *preorganization* to improve binding affinity/kinetics does not sacrifice conformational *flexibility* to accommodate the structural dynamics of the target DNA.

To liberalize reagent access even further, it is worth exploring the minimum number of γ modifications (of any kind) required to improve the editing efficacy relative to the base PNA sequence. Should the ^ser^γ and ^mp^γ modifications prove superior to others reported, such studies, coupled with the directionality^30^ of helical induction, might significantly reduce PNA reagent costs by directing users towards fewer modifications. However, while much of this work should continue, we believe that therapeutic utility of this editing modality will be made more likely by a combination of optimization efforts on all three components (polymer/PNA/donor DNA) of the NP reagents.

## EXPERIMENTAL PROCEDURES

### Resource Availability

#### Lead Contact

Peter M. Glazer

#### Materials Availability

This study did not generate new unique materials.

#### Data and Code Availability

This study did not generate/analyze datasets/code.

### PNAs and DNAs

The sequences for all PNA and γPNA oligomers used in this study targeting the human beta-globin gene were previously published^7^, and are presented in Table 1 with the addition of a sequence containing hydroxymethyl γ-modified bases (^ser^γPNA). All hydroxymethyl-γPNA monomers were purchased from ASM Research Chemicals GmbH (Hannover, Germany), and featured tert-butyloxycarbonyl (boc) and benzyl (Bn) moieties as the protecting groups on the backbone amino and side-chain hydroxyl groups, respectively. The unmodified tcPNA oligomer (designated henceforth as PNA) and the mp-γ-modified tcPNA (^mp^γPNA) were obtained from TruCode Gene Repair, Inc. (San Francisco, CA). We also included ^ser^γPNA-scr (Table 1) as a scrambled sequence control. DNA oligomers utilized for binding and/or melting experiments were purchased from Integrated DNA Technologies (idtdna.com) as either unmodified or 5’-biotinylated derivatives, and their sequences are presented in the relevant Method sections. 50:50 Poly(DL-lactide-co-glycolide), ester terminated with an inherent viscosity 0.55-0.75 (dL/g), was purchased from LACTEL Absorbable polymers (Birmingham, AL). Poly(vinyl alcohol) (PVA), average molecular weight 30,000-70,000 was purchased from Sigma Aldrich (St. Louis, MO). Dichloromethane was purchased from Sigma Aldrich (St. Louis, MO). To test for gene editing by ^mp^γPNA and ^ser^γPNA, we designed 60-nt single-stranded donor DNAs to either introduce the IVS2-654 mutation or to correct it. The sequences for donor DNAs are provided below, with the intended sequence change highlighted in bold. All DNA donors contain three phosphorothioate internucleoside linkages at each end to protect from nuclease degradation.

**Table 1:**
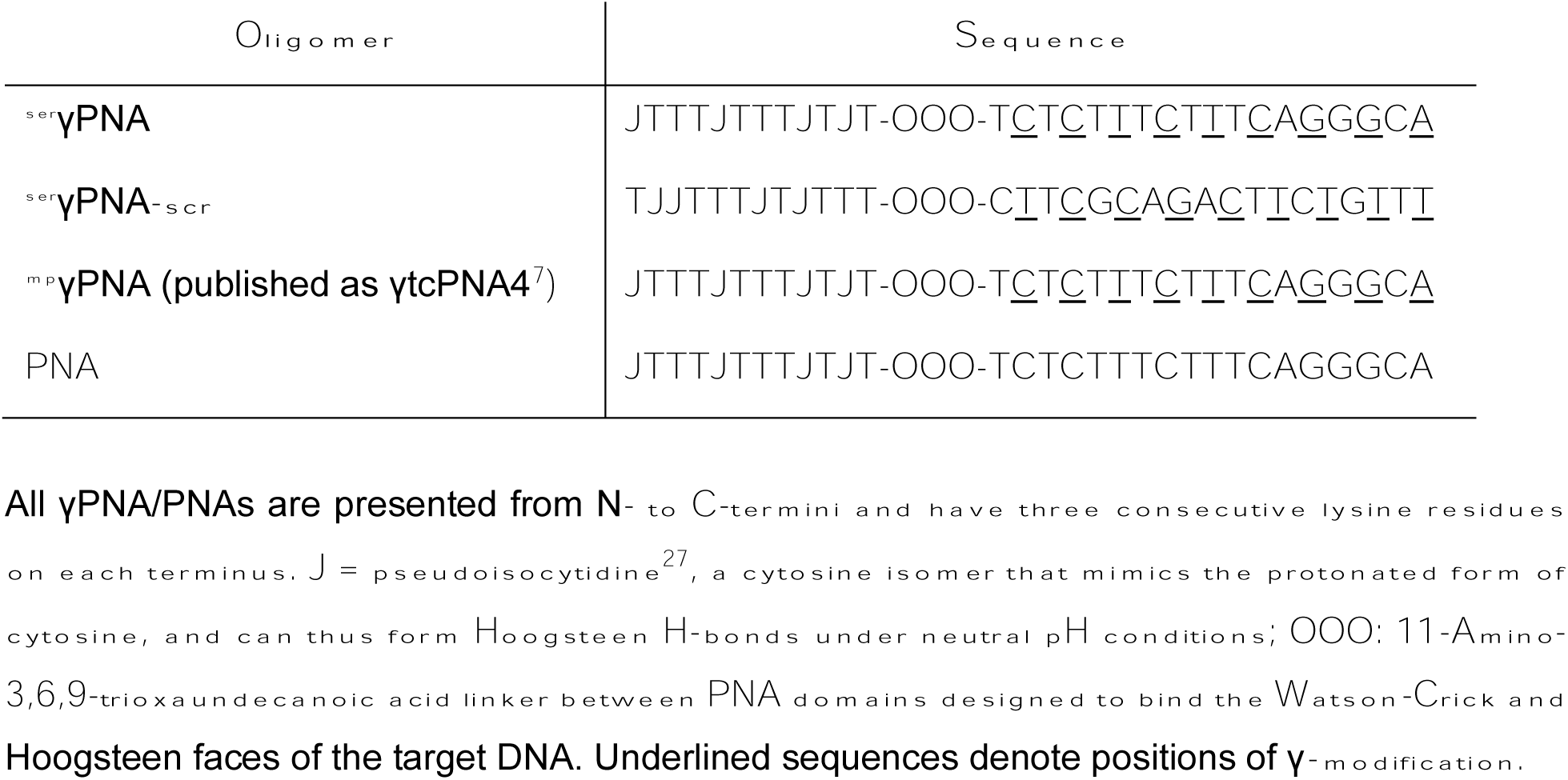
Sequences of ^ser^γPNA, ^ser^γPNA-scr, ^mp^γPNA, and PNA oligomers.

**Correcting donor** (5’-3’): AAAGAATAACAGTGATAATTTCTGGGTTAAGG**C**AATAGCAATATCTCTGCATATAAATAT

**Mutating donor** (5’-3’): AAAGAATAACAGTGATAATTTCTGGGTTAAGG**T**AATAGCAATATCTCTGCATATAAATAT

### PNA Synthesis, Purification, and Characterization

Oligomer synthesis was performed using previously established protocols for standard solid-phase PNA synthesis^51^. Synthesis began on an MBHA solid support (resin) labelled in-house with a lysine residue. The entire synthesis consisted of iterative cycles of two major steps: (1) amine deprotection with a solution of trifluoroacetic acid (TFA)/m-cresol (95/5); and (2) monomer coupling with a cocktail of boc/Bn-protected monomers (A, G, T, C or J), DIEA, and HBTU (1/1.3/0.9) dissolved in *N*-methyl-2-pyrrolidinone and dimethylformamide (1:1). After completion of the cycle for the last monomer, the oligomer was cleaved (released) by submerging the resin in a solution of m-cresol/thioanisole/trifluoroacetic acid/trifluroromethane sulfonic acid (1:1:2:6). The pure product was isolated by reverse phase (RP)-HPLC, using a solvent gradient of acetonitrile and water, and analyzed using MALDI-TOF mass spectrometry.

### Circular Dichroism (CD) Spectropolarimetry

Samples containing increasing concentrations (1-20 µM) of PNA or γPNA oligomers were prepared in a buffer containing 10 mM Na_3_PO_4_ (NaPi, pH 7.4). For the hybrid complexes, samples contained 1 µM complementary ssDNA and 2 µM PNA/γPNA, or 2.5 µM dsDNA and 5 µM PNA/γPNA and were annealed in the same NaPi buffer. For either sample set, a ‘blank’ sample containing the buffer alone was used as a negative control. The annealing step involved high-temperature (95 °C) heating followed by slow cooling to ambient temperature on a heat block to allow formation of the most stable conformations or secondary structures within/by each PNA/γPNA oligomer, or by the respective hybrid complexes. CD spectra were recorded on a Chirascan CD spectropolarimeter (Applied Photophysics). All spectra were collected from 200-350 nm, baseline corrected, and recorded as the average of three consecutive scans. The sequences for the DNA oligos used as ssDNA and dsDNA (ssDNA + ssDNA’) targets are presented below: **ssDNA** (5’-3’; binding sequence underlined): GGTGCAAAGAGGCATGATACATTGTATCATTAT<u>TGCCCTGAAAGAAAGAGA</u>TTAGGGAAAG TATTAGAAATAAGATAAAC **ssDNA’** (5’-3’): GTTTATCTTATTTCTAATACTTTCCCTAATCTCTTTCTTTCAGGGCAATAATGATACAATGTATCATGCCTCTTTGCACC

### Thermal Denaturation Analyses

Melting curves were generated by recording ultraviolet (UV) absorbance at 265 nm with increasing temperature on a Chirascan CD spectropolarimeter equipped with a thermoelectrically regulated multicell holder. All samples were prepared by annealing solutions containing 2.5 µM ss/ds-DNA and 5 µM PNA/γPNA in 10 or 100 mM NaPi buffer (pH 7.4). Absorbance measurements were recorded after every 1 °C change while applying a temperature ramp rate of 1 °C/minute. Where possible, van’t Hoff analyses of the melting curves were performed using the protocols developed by Marky and Breslauer^49^. Briefly, the UV melting curves were transformed into alpha (α) plots to denote the fraction of single strands in hybridized form at each measured temperature. The temperature at α = 0.5 is the melting temperature (Tm) of the hybrid (Table 2).

**Table 2:** T**h**ermodynamic **characterization of PNA/γPNA hybrids with dsDNA target.**

| **(A) 10 mM NaPi<sup>a</sup>** |  |  |  |  |
| --- | --- | --- | --- | --- |
| **Hybrid** | **$-\Delta G$ (kcal mol<sup>-1</sup>)** | **$-\Delta H$ (kcal mol<sup>-1</sup>)** | **$-\Delta S$ (cal mol<sup>-1</sup> K<sup>-1</sup>)** | **T<sub>m</sub> (°C)** |
| PNA/DNA | 35 ± 3 | 77 ± 8 | 141 ± 20 | 52 ± 1 |
| <sup>mp</sup>γPNA/DNA | 34 ± 4 | 69 ± 5 | 118 ± 11 | 52 ± 1 |
| <sup>ser</sup>γPNA/DNA | 34 ± 2 | 74 ± 8 | 134 ± 13 | 50 ± 1 |
| **(B) 100 mM NaPi<sup>a</sup>** |  |  |  |  |
|  | **$-\Delta G$ (kcal mol<sup>-1</sup>)** | **$-\Delta H$ (kcal mol<sup>-1</sup>)** | **$-\Delta S$ (cal mol<sup>-1</sup> K<sup>-1</sup>)** | **T<sub>m</sub> (°C)** |
| PNA/DNA | 48 ± 3 | 139 ± 11 | 306 ± 16 | 73 ± 2 |
| <sup>mp</sup>γPNA/DNA | 41 ± 6 | 94 ± 8 | 176 ± 10 | 72 ± 1 |
| <sup>ser</sup>γPNA/DNA | 45 ± 5 | 124 ± 9 | 264 ± 11 | 72 ± 1 |
| <sup>a</sup>All values are calculated for 25 °C. |  |  |  |  |

### Electrophoretic Mobility Shift (Gel-Shift) Assays

Strand invasion by each PNA/γPNA was evaluated using two different duplex substrates. First, 50 nM of a 100mer dsDNA target, preassembled by hybridizing ssDNA with its complement (ssDNA’), was annealed with increasing equivalents (0 – 10-fold) of PNA/γPNA to assemble the most stable hybrid. Each sample was then run on a nondenaturing 8 % polyacrylamide gel electrophoresis (PAGE) system in 1x tris borate EDTA (TBE) buffer (pH 7.4) at 120 V for 40 min. The gel was stained with 1x SYBR Gold (Invitrogen/ThermoFisher Scientific, catalog #S11494) for 5 min and washed with 1x TBE for 20 min. Resolved bands for free and bound DNA were visualized on a Biorad Universal Hood II gel doc system, and the images were processed and inverted using Image Lab software (version 5.2.1, Biorad).

### Nanoparticle Formulation

PLGA NPs encapsulating PNA, and donor DNA were formulated using a double-emulsion solvent evaporation technique. Briefly, 40 mg of PLGA was dissolved in 1 mL of dichloromethane (40 mg ml^-1^). Forty μl of donor DNA (1 mM) and 80 μl of PNA (1 mM) were quickly mixed and added to the polymer solution dropwise while vortexing the polymer. The mixture was subsequently sonicated at 38% amplitude 3 times for 10 seconds using a probe sonicator. To form the second emulsion, the primary emulsion was added dropwise to 2 mL of a 5% (w/v) solution of PVA. The second emulsion was subsequently sonicated at 38% amplitude 3 times for 10 seconds using a probe sonicator. This mixture was finally poured into 20 mL of 0.3% (w/v) PVA solution and stirred for 3 hours at room temperature (360 rpm) while the NPs ‘hardened.’ NPs were subsequently pelleted by centrifugation at 16,100 g for 15 minutes. The NP pellet was resuspended in 20 mL of diH_2_O and washed an additional 2 times, for a total of 3 centrifugation steps. The final NP pellet was resuspended in diH_2_O with trehalose (5 mg/ml) such that the final ratio of NP:trehalose was 1 mg NP:1 mg trehalose. The final NP solution was flash frozen in liquid nitrogen, lyophilized for 48 hours, and stored at −20°C until further use.

### Nanoparticle Characterization

NP size and zeta potential were measured using the Malvern ZetaSizer as per manufacturer guidelines. Measurements were made in diH_2_O at a NP concentration of 0.05 mg/ml. Total nucleic acid loading was determined by absorbance at 260 nm following de-formulation in DMSO. Total nucleic acid release was determined by incubating 2 mg of NPs in 1 mL of 1X PBS in a 37°C shaker. At specified timepoints, NPs were centrifuged at 21000 g; 950 μl of the supernatant was collected and replaced, followed by absorbance measurement at 260 nm.

### *Ex vivo* Gene Editing in Primary Bone Marrow Cells

Bone marrow cells were harvested by flushing the femurs and tibias of Townes mice containing the human β-globin transgene. 500,000 cells were subsequently treated with 2 mg ml^-1^ PLGA NPs in Roswell Park Memorial Institute (RPMI) medium supplemented with 20% FBS and 5% penn-strep. 72 hours later, genomic DNA (gDNA) was harvested using the Promega ReliaPrep^TM^ gDNA Tissue Miniprep System (Promega, Madison, WI). Similarly, bone marrow cells from a mouse model of β-thalassemia with the IVS2-654 mutation^52^ were flushed and treated with PLGA NPs containing PNA and a ‘correcting donor’ DNA. As before, gDNA was harvested 72 hours after NP treatment. DNA size selection was performed using the Ampure beads (1.8x) following gDNA extraction when comparing ^ser^γPNA to ^ser^γPNA-scr. In all cases, editing frequencies were quantified using a droplet digital (dd) PCR method which we previously developed and validated^8^. Each ddPCR reaction consisted of up to 80 ng of gDNA, 12.5 μl of 2 × ddPCR™ supermix for probes (no dUTP) (Bio-Rad, Hercules, CA), 0.225 μl forward primer (100 μM), 0.225 μl reverse primer (100 μM), 0.063 μl β-thal probe (100 μM), 0.063 μl wild-type probe (100 μM) (Integrated DNA Technologies, Coralville, IA), 0.5 μl EcoR1, 11.424 μl gDNA and dH_2_O. Droplets were generated using the Automated Droplet Generator (AutoDG™, Bio-Rad). Thermocycling conditions were as follows: 95 °C 10 min, (94 °C 30 s, 55.3 °C 5 min – ramp 2 °C/s) x 39 cycles, 98 °C 10 min, hold at 4 °C. Droplets were allowed to rest at 4 °C for at least 30 min after cycling and were then read using the QX200 ™ Droplet Reader (Bio-Rad). Data were analyzed using QuantaSoft™ software. Data are represented as the fractional abundance of the wild-type allele. The primers used for ddPCR were as follows: forward (5’-3’): ACCATTCTAAAGAATAACAGTGA, reverse (5’-3’): CCTCTTACATCAGTTACAATTT. The probes used for ddPCR were as follows: wild-type (5’-FAM): TGGGTTAAGG**C**AATAGCAA; β-thal (5’-HEX): TCTGGGTTAAGG**T**AATAGCAAT, where the base in bold is complementary to the targeted mutation site.

## ACKNOWLEDGMENTS.

This work was supported by R01HL139756 and U01AI145965 (to PMG and WMS), R35CA197574 (to P.M.G.) and the NIGMS Medical Scientist Training Program T32-GM07205 (to E.Q.).

## AUTHOR CONTRIBUTIONS

SNO, EQ, JDRP, and HKM performed experiments. WMS and PMG supervised the work and analyzed data. SNO, EQ, RB, JDRP, and PMG wrote the manuscript.

## DECLARATION OF INTERESTS

PMG is a consultant to and has equity in Gennao Bio, Cybrexa Therapeutics, and pHLIP Inc., none of which are related to the work reported here. PMG, WMS, EQ, and RB are inventors on patents assigned to Yale University related to gene editing by PNAs.

## SUPPLEMENTAL INFORMATION

Document S1. Supplemental Figures S1–S5

**Figure S1:**
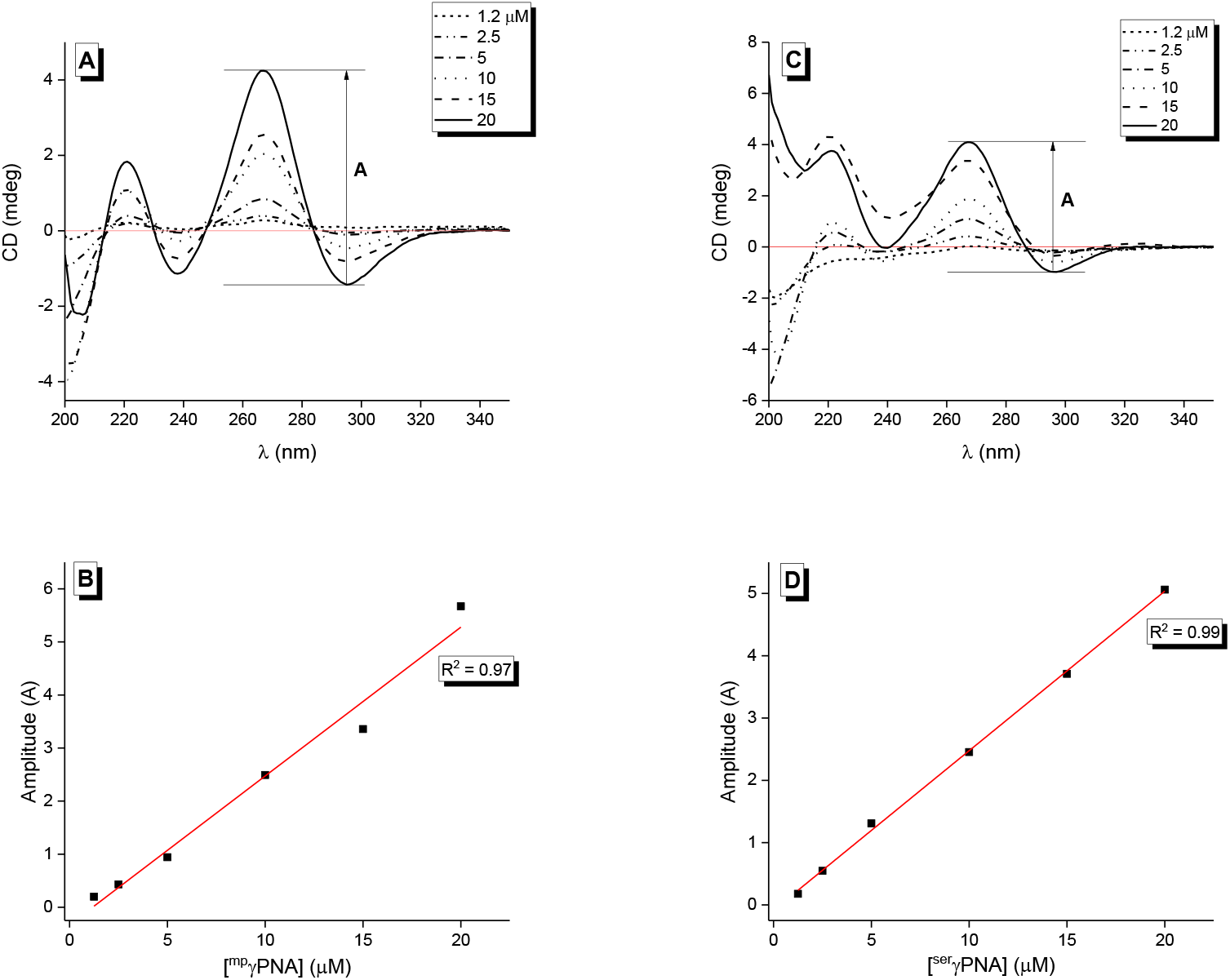
CD spectra of ^mp^γPNA and ^ser^γPNA over concentrations ranging from 1.2 – 20 µM. (A) Concentration dependent CD spectra for ^mp^γPNA; (B) Plot of amplitude against oligomer concentration for ^mp^γPNA; (C) Concentration dependent CD spectra for ^ser^γPNA; (D) Plot of amplitude against oligomer concentration for ser?PNA. All samples were prepared in 10 mM NaPi buffer and spectra recorded at 37°C.

**Figure S2:**
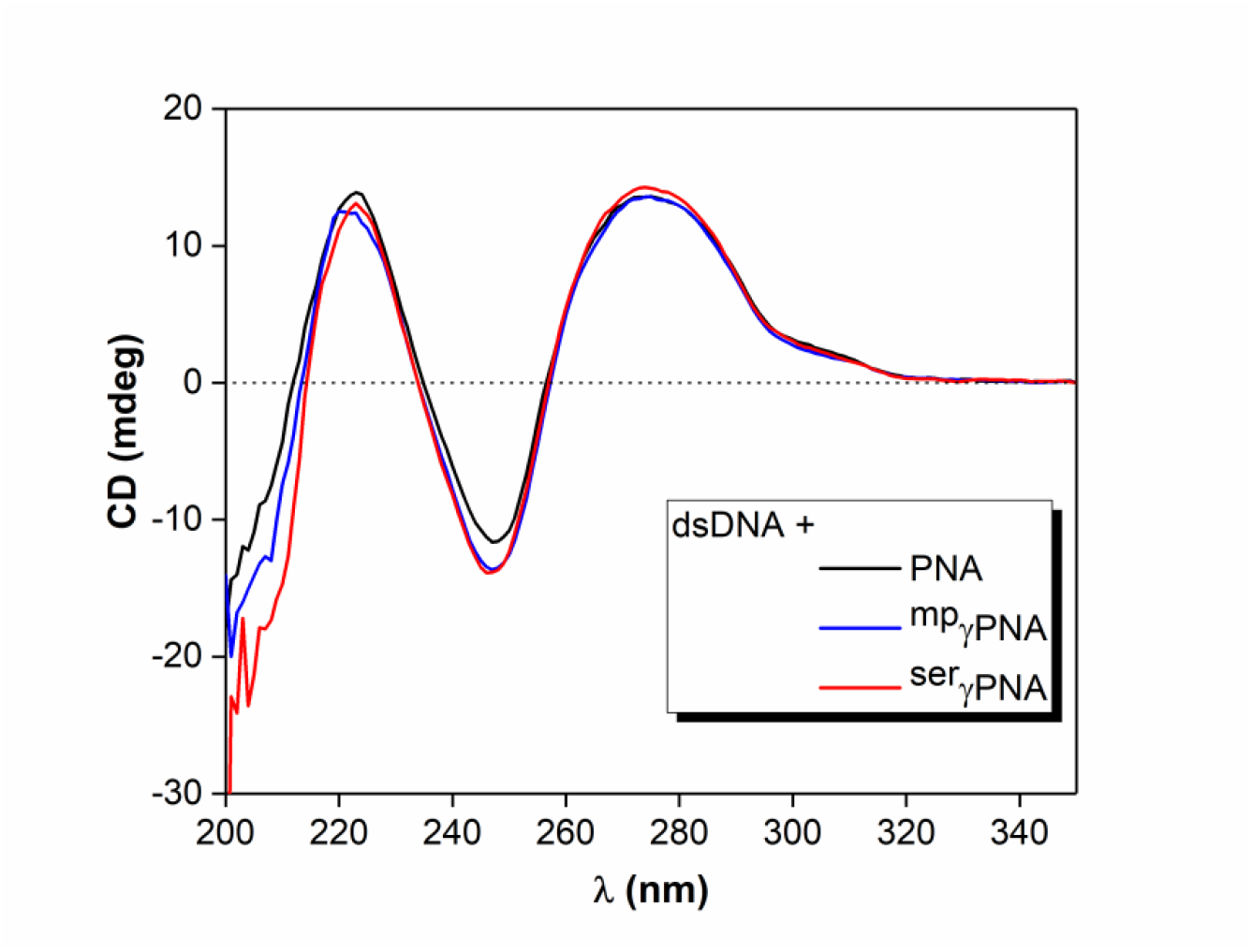
CD spectra of dsDNA-PNA/γPNA hybrids. Samples contained 2.5 μM dsDNA alone or in combination with 5 μM PNA (black), ^mp^γPNA (blue), or ^ser^γPNA (red), and were annealed in 10 mM NaPi buffer as in Methods.

**Figure S3:**
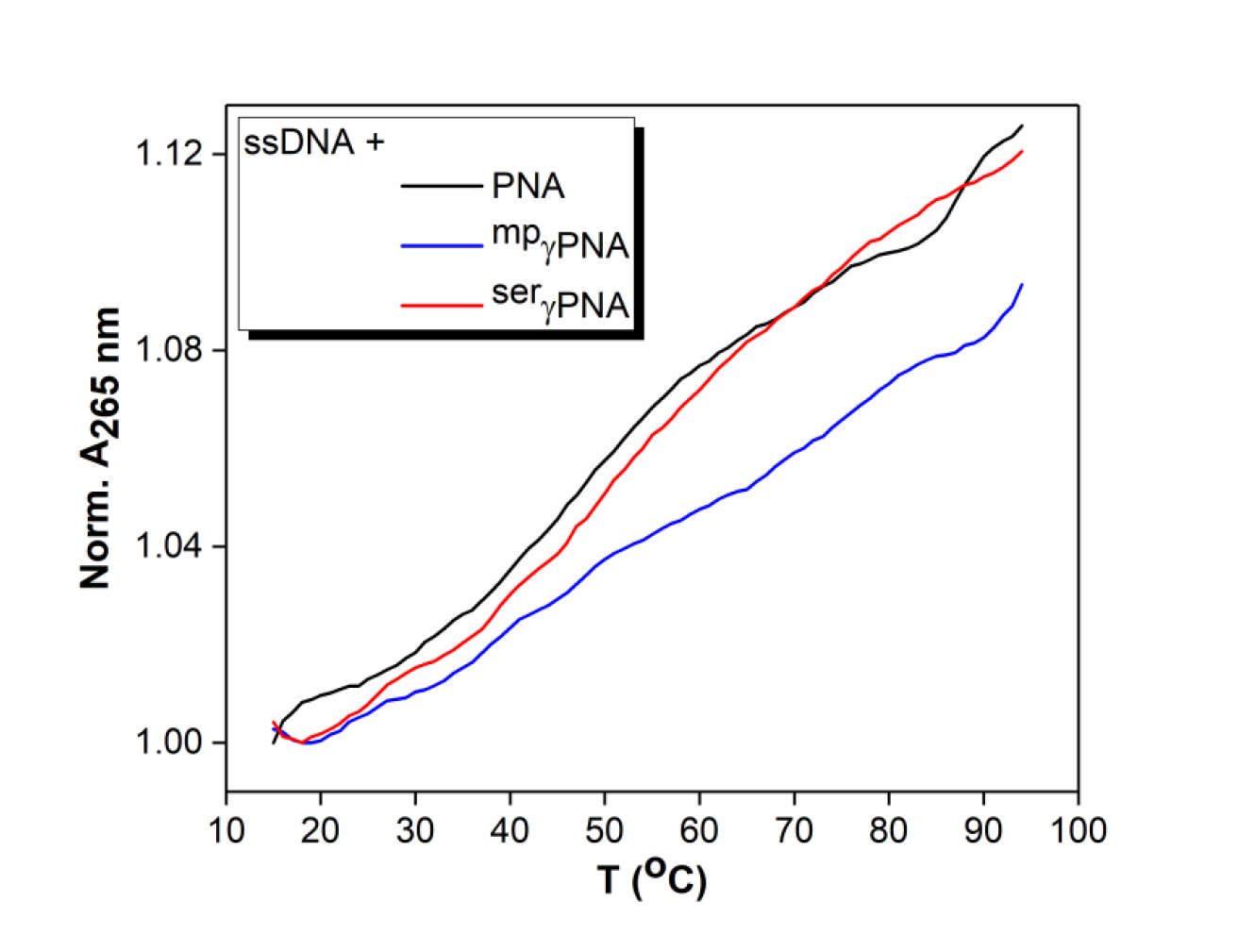
UV-thermal denaturation analyses on ssDNA-PNA/γPNA hybrids. Samples contained 2.5 μM dsDNA alone or in combination with 5 μM PNA (black), ^mp^γPNA (blue), or ^ser^γPNA (red), and were annealed in 10 mM NaPi buffer as in Methods.

**Figure S4:**
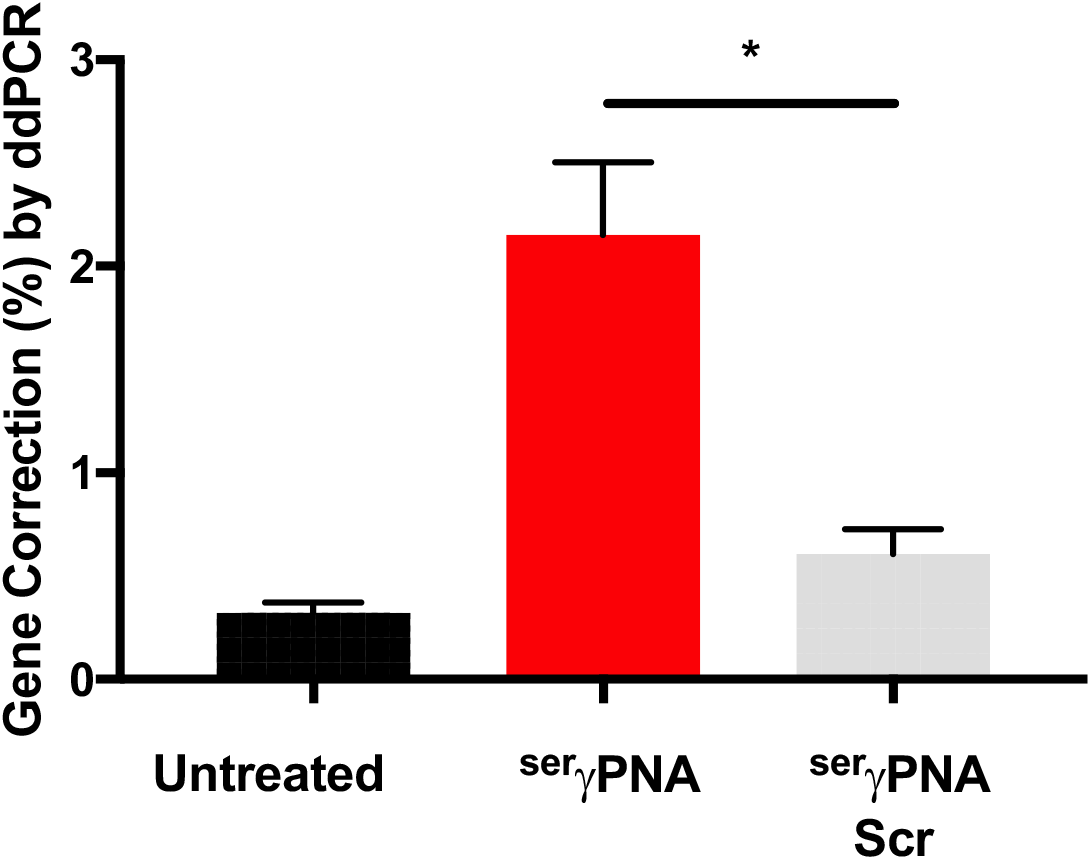
Correction of the β-thalassemia causing mutation in primary bone marrow cells. BMCs were left untreated (black) or were treated with NPs containing ^ser^γPNA (red), or ^ser^PNA-scr (gray).

